# Interferon inhibits a model RNA virus via a limited set of inducible effector genes

**DOI:** 10.1101/2023.02.21.529297

**Authors:** Matthew B. McDougal, Anthony M. De Maria, Maikke B. Ohlson, Ashwani Kumar, Chao Xing, John W. Schoggins

## Abstract

Interferons control viral infection by inducing the expression of antiviral effector proteins encoded by interferon-stimulated genes (ISGs). The field has mostly focused on identifying individual antiviral ISG effectors and defining their mechanisms of action. However, fundamental gaps in knowledge about the interferon response remain. For example, it is not known how many ISGs are required to protect cells from a particular virus, though it is theorized that numerous ISGs act in concert to achieve viral inhibition. Here, we used CRISPR-based loss-of-function screens to identify a markedly limited set of ISGs that confer interferon-mediated suppression of a model alphavirus, Venezuelan equine encephalitis virus (VEEV). We show via combinatorial gene targeting that three antiviral effectors – ZAP, IFIT3, and IFIT1 – together constitute the majority of interferon-mediated restriction of VEEV, while accounting for less than 0.5% of the interferon-induced transcriptome. Together, our data suggests a refined model of the antiviral interferon response in which a small subset of “dominant” ISGs may confer the bulk of the inhibition of a given virus.

## Introduction

Interferon (IFN) is a cytokine that exerts antiviral properties by inducing the expression of hundreds of interferon-stimulated genes (ISGs), some of which encode effector proteins with specific mechanisms of action that restrict viral infection (Schoggins, 2019). IFN is broadly antiviral and inhibits diverse viruses with both RNA and DNA genomes (Isaacs & Lindenmann, 1957; Isaacs *et al*, 1957). It has been theorized that IFN exhibits these broad antiviral properties because many ISG effectors act in concert to suppress viral infection at multiple steps of the viral replication cycle (i.e. entry, uncoating, translation, replication, assembly, egress) (Jones *et al*, 2021; Schneider *et al*, 2014).

Major goals in the ISG field have focused on answering key questions. The first question is “which ISGs have antiviral activity?” This has primarily been addressed through genetic screens. Over the last 10+ years, we and others have used cDNA-based overexpression screens to identify ISGs that restrict diverse viruses (Bamford *et al*, 2022; Boys *et al*, 2020; Dittmann *et al*, 2015; Feng *et al*, 2018; Hanners *et al*, 2021; Itsui *et al*, 2006; Kane *et al*, 2016; Kuroda *et al*, 2020; Liu *et al*, 2012; Pfaender *et al*, 2020; Qi *et al*, 2017; Rihn *et al*, 2019; Schoggins *et al*, 2014; Schoggins *et al*, 2011; Wickenhagen *et al*, 2021; Wilson *et al*, 2012; Zhang *et al*, 2007). Overexpression screening is a powerful approach, but it has disadvantages. Expressing a single gene only allows for the identification of effectors that are sufficient to inhibit viral infection, but not those that are required for IFN-mediated suppression. Overexpressing individual genes may also miss effectors that work in multi-protein complexes to exert antiviral functions. Recently, CRISPR-Cas9 gene silencing screens have begun to address some of these limitations. By genetically silencing individual genes in the context of a functional IFN response, CRISPR screens have the potential to identify those ISGs that are necessary for IFN-mediated protective effects (Dang *et al*, 2022; Mac Kain *et al*, 2022; OhAinle *et al*, 2018; Richardson *et al*, 2018). A second question of interest in the field is “what are the mechanisms of ISG effectors?” Mechanistic studies include determining which stage in the viral replication cycle an ISG effector acts and the molecular details of how the effector achieves viral inhibition. A logical third question is “what is the physiological role of an ISG effector during viral infection?” The clearest answers to this question are frequently obtained from pathogenesis studies using ISG knockout mice (Schoggins, 2014).

Beyond these traditional goals of identifying and characterizing ISGs, there are other more esoteric and theoretical questions about ISGs that have yet to be answered. These include, “how many ISGs are required to inhibit a given virus?” “Do ISGs work in concert with each other?” “Are some ISGs more potent than others?” “Do different ISGs have varying degrees of antiviral ‘value’?” Remarkably, some of these questions were acknowledged decades before the current era of high-throughput genetic screens. In their 1986 study on the anti-influenza potential of MX1, when only a total of 12 ISGs were known at that time, Staeheli et al discussed the IFN response and presciently wrote, “it is not known which or how many of these proteins are required for protection against a particular virus” (Staeheli *et al*, 1986). In the 35+ years following this study, the question of “which” ISGs are antiviral has been a key driver in the field. However, the question of “how many” ISGs are required for protection remains obscure. In this study, we present an ISG screening workflow that yielded surprising results, leading us to examine some of these more enigmatic concepts of ISG biology.

Recent CRISPR-based ISG discovery screens have focused on yellow fever virus (Richardson *et al*., 2018), HIV-1 (OhAinle *et al*., 2018), and SARS-CoV-2 (Mac Kain *et al*., 2022). We sought to use similar approaches to identify potentially novel ISGs that inhibit alphaviruses. These are positive-sense single-stranded RNA viruses in the *Togaviridae* family, which includes several model and pathogenic viruses. Using Venezuelan equine encephalitis virus (VEEV) as a representative alphavirus in a CRISPR/ISG screening workflow, we found that a surprisingly small number of known effectors inhibit this virus. Combinatorial ISG silencing demonstrated that the majority of the IFN-mediated suppression of this virus was dominated by just three ISGs out of more than 600 induced transcripts. Furthermore, these three ISGs did not dominate the IFN suppression of certain related and unrelated viruses, suggesting that a distinct set of dominant antiviral effectors are required for the restriction of different viruses. Other reported anti-alphavirus ISGs did not contribute to IFN-mediated restriction of VEEV in the absence of the “dominant” ISGs we identified here. We propose a model in which a limited set of dominant ISGs make major contributions to IFN-mediated viral restriction in a virus-specific manner.

## Results

To identify the genes required for IFN-mediated restriction of a GFP-expressing VEEV (VEEV-GFP), we performed genome-wide and ISG-targeted CRISPR-Cas9 knockout screens. U-2 OS cells were transduced with a pooled Brunello lentivirus library to target all human genes (Doench *et al*, 2016), or with a pooled ISG-specific library targeting approximately 1,900 genes (Roesch *et al*, 2018). CRISPR-targeted cells were treated with a dose of IFNβ that is nearly completely suppressive in normal cells, and then infected with VEEV-GFP. CRISPR-targeted cells that became infected despite the presence of IFN were FACS-sorted and sgRNAs were PCR-amplified and sequenced (Fig 1A, Table S1, S2). The screens identified all genes involved in JAK/STAT signaling (FDR <0.25), confirming the dependence of viral suppression on IFN signaling (Fig 1B). Additionally, previously characterized anti-VEEV effectors IFIT1 and IFIT3 were identified in both screens (Fig 1B, 1C), underscoring the robustness and accuracy of the data.

**Figure 1.**
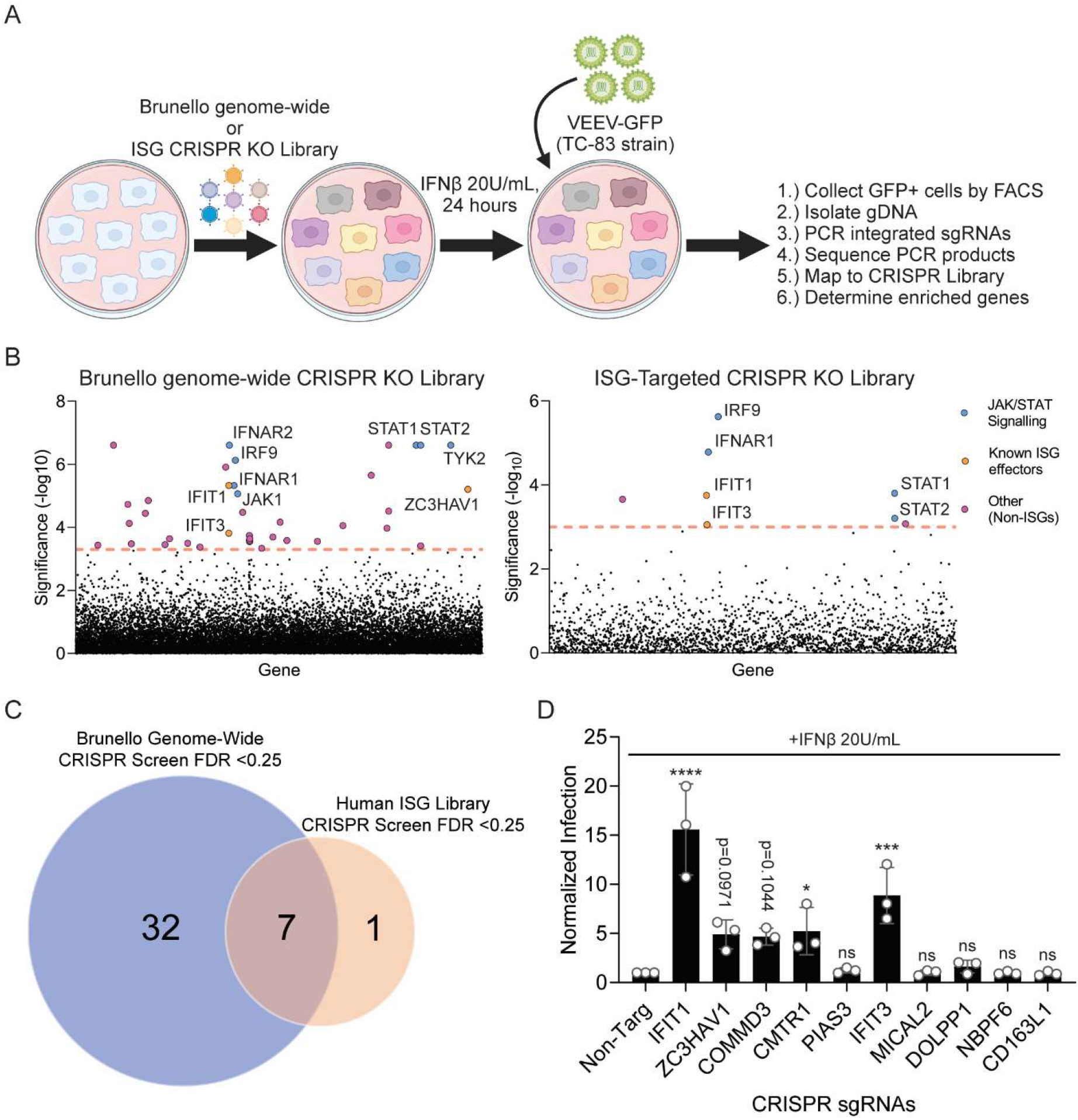
CRISPR Screening identifies a limited set of genes that restrict VEEV infection. A. Schematic of genome-wide and ISG-targeted CRISPR screen to identify genes required for IFN-mediated suppression of VEEV. Graphic generated with Biorender. B. Manhattan dot plots of genome-wide CRISPR screen (left) or ISG-targeted CRISPR screen (right) results. Significance was determined by MAGeCK analysis, and the dotted lines represent a cut-off of FDR<0.25. Genes are plotted in alphabetical order. C. Venn diagram comparing the CRISPR screening hits (FDR<0.25) from the genome-wide (blue) and ISG-targeted (orange) screens. D. Validation of selected CRISPR screening hits by targeted gene silencing after IFN treatment. Genes are plotted on the x-axis from lowest FDR value (IFIT1) to highest FDR value (CD163L1) as determined by MAGeCK analysis in the genome-wide screen. All samples were normalized to non-targeting controls (“Non-Targ”) for graphing. One-way ANOVA with Dunnett’s test comparing each group to the non-targeting control was performed on non-normalized data, n=3.

We next selected 10 genes, both known ISGs and non-ISGs, to validate the CRISPR screening hits. Each gene was targeted by two independent sgRNAs, expressed from either a puromycin- or blasticidin-selectable lentiCRISPRv2 vector. Pooled cell populations transduced with both lentiviruses were selected in puromycin and blasticidin. Selected cells were treated with IFN, followed by infection with VEEV-GFP. This approach validated the known anti-VEEV ISGs *IFIT1*, *IFIT3*, and *ZC3HAV1*, and *CMTR1* which was recently shown to control the expression of certain ISG-encoded proteins (Williams *et al*, 2020) (Fig 1D). We also found that cells targeted for silencing of *COMMD3*, a gene linked to endosomal recycling and chemotaxis of lymphocytes (Maine & Burstein, 2007; Nakai *et al*, 2019; Singla *et al*, 2019), were modestly less susceptible to IFN-mediated restriction of VEEV. In addition to *COMMD3*, other genes encoding components of the CCC (<u>C</u>OMMD/<u>C</u>CDC22/<u>C</u>CDC93) complex (*CCDC22, CCDC93*) were hits in the genome-wide CRISPR Screen (Table S1, S2). The CCC complex also interacts with the WASH (<u>W</u>iskott–<u>A</u>ldrich syndrome protein and <u>S</u>CAR <u>h</u>omologue) complex and Retriever complex for performing functions involved in endosomal trafficking (Chen *et al*, 2019; Wang *et al*, 2018). Components of these complexes including *VPS35L/C16orf62, KIAA1033/WASHC4*, and *FAM21C/WASHC2C*) were also hits in the genome-wide CRISPR screen, but were not validated here (Table S1, S2). Other genes tested during these validation experiments, most of which were between 0.1 and 0.25 FDR in the initial screen, did not validate (Fig 1D). Together, the screens and validation assays identified a unique set of genes required for IFN-mediated restriction of VEEV infection.

To contextualize the CRISPR screening data relative to the IFN response, we determined which genes were upregulated in U-2 OS cells after IFNβ treatment. We performed RNA-Seq after treating cells for 6 and 24 hours with the same dose of IFNβ as used in the screens (Fig 2A, 2B). We identified 620 genes significantly induced after IFNβ treatment (log_2_FC>1 and p_adj_<0.5 at 6- or 24-hour treatment) (Fig 2C). Of the 40 genes revealed as hits in the CRISPR screens, only 6 (*STAT1*, *STAT2*, *IRF9*, *ZC3HAV1*, *IFIT3*, and *IFIT1*) were significantly induced by IFN in the RNA-Seq dataset (Fig 2C, 2D). Three of these genes (*STAT1*, *STAT2*, and *IRF9*) participate in the JAK/STAT signaling cascade and are themselves ISGs but do not have known direct antiviral effector functions. The other three genes, *ZC3HAV1*, *IFIT3*, and *IFIT1* encode known antiviral effectors that inhibit viral translation. ZAP binds to CpG dinucleotides present in alphavirus RNA (Bick *et al*, 2003; Guo *et al*, 2004; Takata *et al*, 2017), while IFIT1 and IFIT3 associate to bind to 5’ ends of viral Cap0 RNA (RNA that is capped but lacking 2’-O-methylation) and inhibit translation (Daffis *et al*, 2010; Hyde *et al*, 2014; Johnson *et al*, 2018). These results suggest a highly prominent role for ZAP, IFIT3, and IFIT1 in suppressing VEEV infection.

**Figure 2.**
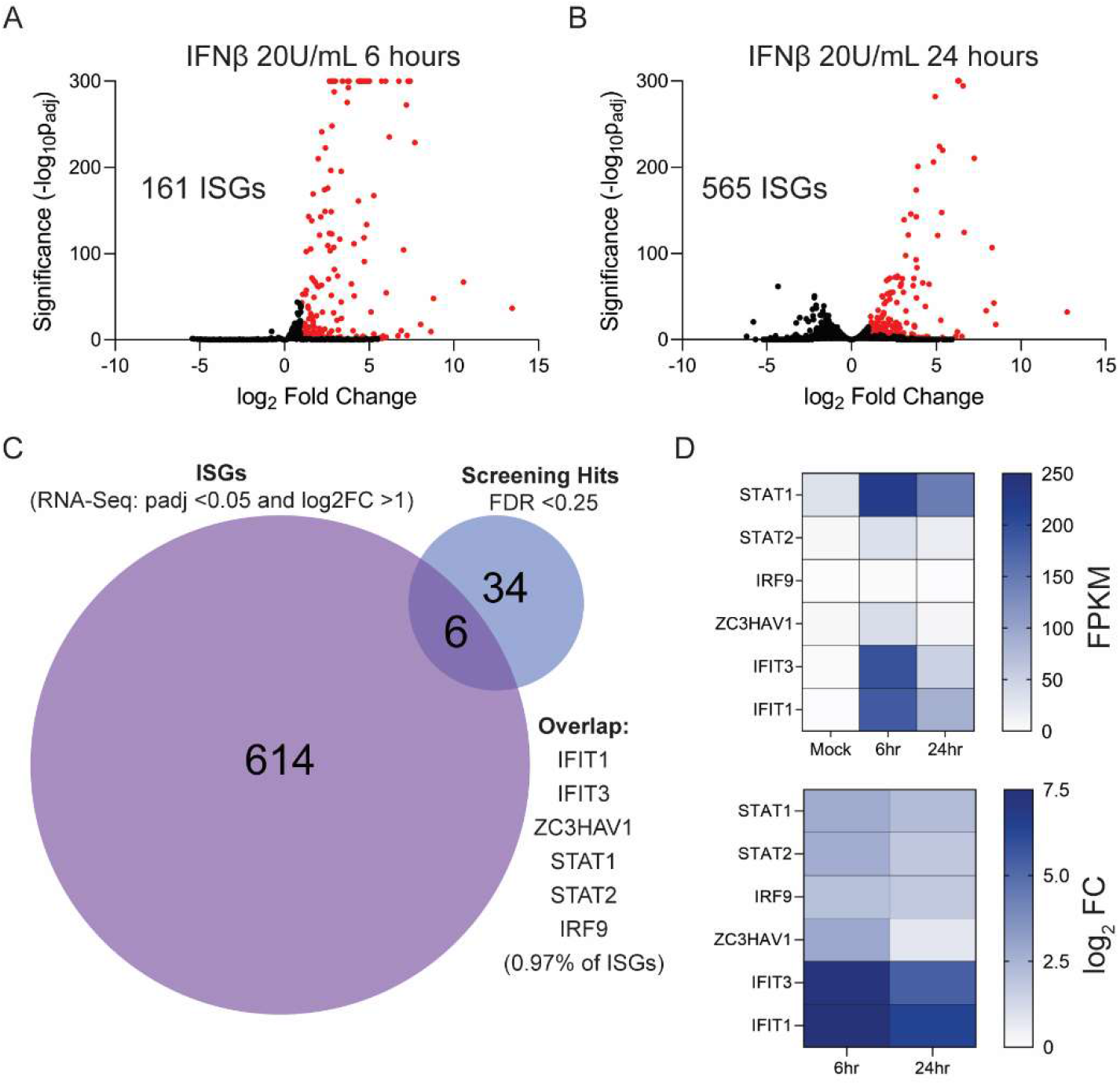
RNA-Seq reveals a limited set of IFN-induced genes were identified as CRISPR screening hits. A. Volcano plot of differentially expressed genes during RNA-Seq of U-2 OS cells treated with IFNβ for 6 hrs. Significance was determined by DESeq2, and red dots represent genes with a log_2_ fold-change > 1 and a p_adj_ < 0.05. B. Volcano plot of differentially expressed genes during RNA-Seq of U-2 OS cells treated with IFNβ for 24 hrs. Significance was determined by DESeq2, and red dots represent genes with a log_2_ fold-change > 1 and a p_adj_ < 0.05. C. Venn diagram of CRISPR screening hits (FDR<0.25) and ISGs as determined by RNA-Seq, with overlapping genes listed. D. Heat maps of FPKM and log_2_FC values (from RNA-Seq) of IFN-induced CRISPR screening hits at 6 and 24 hrs post IFN-treatment.

By coupling CRISPR screening with RNA-Seq (Fig 1 and Fig 2), we identified only 3 genes out of 620 IFN-induced transcripts (0.48% of IFN-induced transcripts) that appear to contribute to IFN-mediated suppression of VEEV. We found this striking, as it is commonly thought that IFN suppresses viruses by the concerted action of numerous ISG-encoded effectors (Chen *et al*, 2021; Feng *et al*., 2018; Hubel *et al*, 2019; Jones *et al*., 2021; Metz *et al*, 2012; Randall & Goodbourn, 2008; Schneider *et al*., 2014). Our data invoke an alternative hypothesis, that ZAP, IFIT3, and IFIT1 may be serving as “dominant” antiviral effectors that are particularly important for IFN-mediated suppression of VEEV infection. To test this, we generated multiplexed CRISPR-targeted cells expressing 3 sgRNAs to silence ZC3HAV1, IFIT3, and IFIT1 (herein referred to as “ZAP/IFIT3/IFIT1-targeted cells” or to silence IRF9, STAT1, and STAT2 (herein referred to as “IRF9/STAT1/STAT2-targeted cells”) as a control. Cells were pre-treated with IFN, followed by infection with VEEV-GFP and quantification of infectivity by flow cytometry. We confirmed by Western blot that the triple CRISPR guide strategy resulted in marked reductions in protein abundance (Fig 3A, 3B, Fig S1A, S1B). As predicted, IFN-treated IRF9/STAT1/STAT2-targeted U-2 OS cells were highly permissive to VEEV-GFP infection when compared to IFN-treated cells expressing non-targeting CRISPR guides (Fig 3A). Notably, ZAP/IFIT3/IFIT1-targeted cells were infected at nearly the same rate as IRF9/STAT1/STAT2-targeted cells (85% versus 73%), although they still exhibited a statistically significant decrease in infectivity (Fig 3A). Similar experiments in ZAP/IFIT3/IFIT1-targeted Huh7.5 cells demonstrated that these three effectors were crucial for restricting VEEV after IFNβ treatment (Fig 3B). These results indicate that the dominant antiviral effects of ZAP, IFIT3, and IFIT1 are likely not specific to a certain cell type.

**Figure 3.**
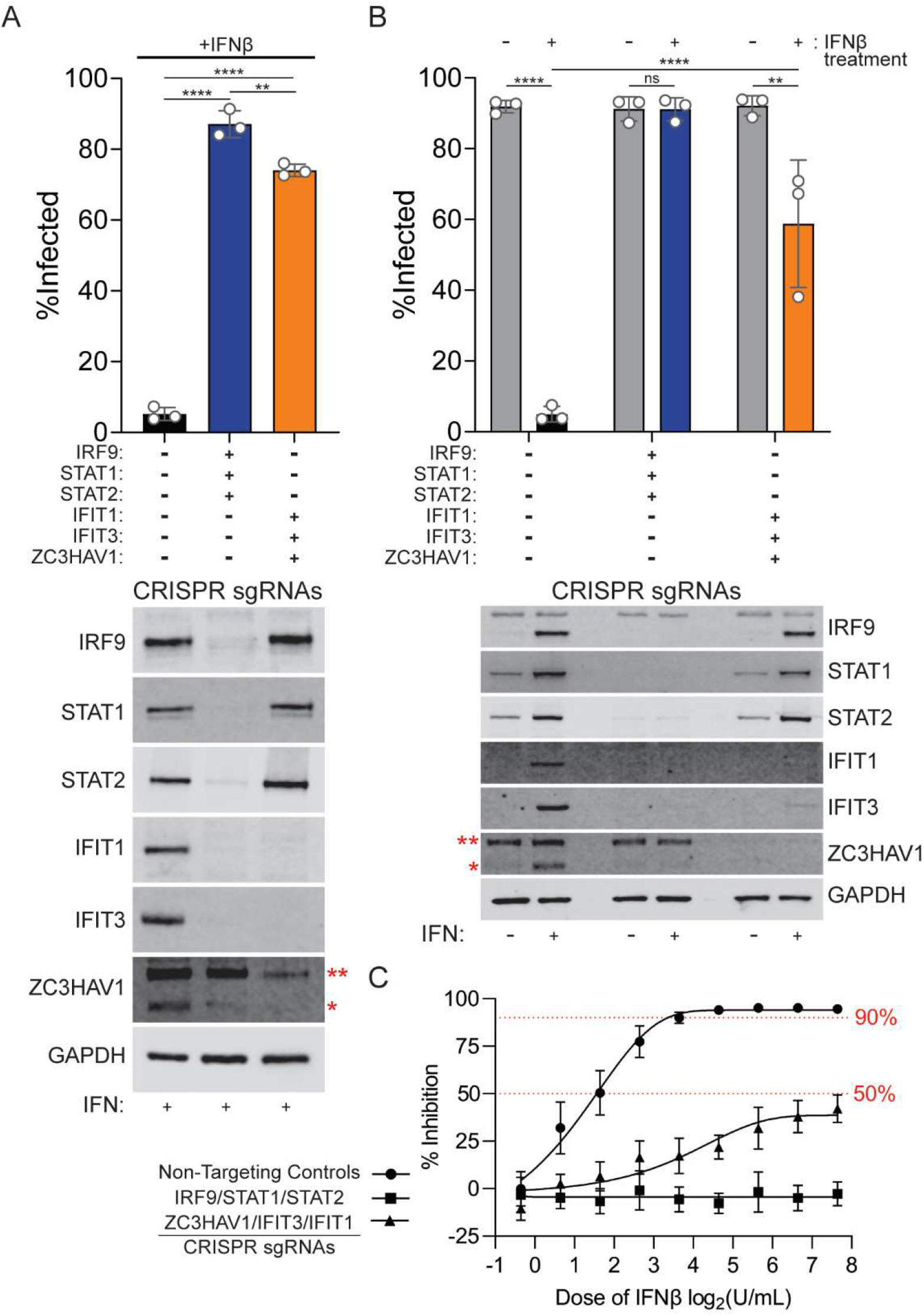
ZAP, IFIT3, and IFIT1 are dominant contributors to the IFN-mediated suppression of VEEV. A. (Top) Effects of CRISPR-based gene silencing of IRF9, STAT1, and STAT2 or ZC3HAV1, IFIT3, and IFIT1 on VEEV infection (MOI of 10 for 5 hrs) in U-2 OS cells with IFNβ pretreatment (20U/mL for 24 hrs). n=3, One-way ANOVA with Dunnett’s test. (Bottom) Western blot showing reduced protein abundance in CRISPR-targeted cells. Red asterisks denote long (**) and short (*) isoforms of ZAP. B. (Top) Effects of CRISPR-based gene silencing of IRF9, STAT1, and STAT2 or ZC3HAV1, IFIT3, and IFIT1 on VEEV infection (MOI of 10 for 4.5 hrs) in Huh7.5 cells with (+ symbols) and without (- symbols) IFNβ pretreatment (100U/mL for 24 hrs). n=3, One-way ANOVA with Dunnett’s test. (Bottom) Western blot showing CRISPR targeting efficiency at the protein-level. Red asterisks denote long (**) and short (*) isoforms of ZAP. C. Dose-curve of IFNβ pretreatment (2-fold dilutions from 200U/mL to 1.5625U/mL for 24 hrs) demonstrating inhibition of VEEV infection (MOI of 5 for 5 hrs) in control (circles), IRF9/STAT1/STAT2-targeted (squares), or ZAP/IFIT3/IFIT1-targeted (triangles) U-2 OS cells.

As one dose of IFNβ had been used for all previous experiments in U-2 OS cells, we sought to determine if high doses of IFNβ treatment could restrict VEEV in ZAP/IFIT3/IFIT1-targeted cells. Cells were treated with 2-fold serial dilutions of IFNβ from 200 U/mL (10-fold higher than the suppressive dose used in all previous experiments) to 1.6 U/mL and infected with VEEV-GFP at an MOI of 5 (Fig 3C). In non-targeting control cells, IFN inhibited VEEV in a dose-dependent manner, with 50% inhibition at 3 U/mL and 90% inhibition at 10 U/mL. All higher doses maximally inhibited infection by 95%. In IRF9/STAT1/STAT2-targeted cells, IFN did not inhibit infection at any dose. In ZAP/IFIT3/IFIT1-targeted cells, we could not achieve 50% inhibition at even the highest dose; maximal inhibition was 42% at 200 U/mL. Although we cannot calculate an IC_50_ for IFN suppression in ZAP/IFIT3/IFIT1-targeted cells based on these data, we infer that if 50% inhibition could be achieved at higher doses, the IC_50_ relative to control cells would increase greater than 60-fold (IC_50_ = 3 U/mL vs IC_50_ > 200 U/mL). These data indicate that over a wide range of IFN doses, ZAP, IFIT3, and IFIT1 together comprise an outsized proportion of the total IFN-mediated suppression of VEEV-GFP.

ZAP, IFIT3, and IFIT1 have been shown to exhibit broad antiviral activity towards diverse viruses (Andrejeva *et al*, 2013; Bick *et al*., 2003; Chikhalya *et al*, 2021; Fensterl & Sen, 2015; Gao *et al*, 2002; Gonzalez-Perez *et al*, 2021; Griffante *et al*, 2021; Hyde *et al*., 2014; Johnson *et al*., 2018; Pichlmair *et al*, 2011; Reynaud *et al*, 2015; Schmeisser *et al*, 2010; Zhang *et al*, 2022). We therefore used gene silencing to determine the contribution of ZAP, IFIT3, and IFIT1 to the antiviral activity of IFN towards other IFN-sensitive viruses. We challenged ZAP/IFIT3/IFIT1-targeted U-2 OS cells with the alphavirus O’nyong ‘nyong virus (ONNV) or the unrelated flavivirus Zika virus (ZIKV) after pretreatment with IFNβ (Fig 4A). Silencing of ZAP, IFIT3, and IFIT1 had little to no effect on IFN-mediated suppression of ONNV and ZIKV. We also challenged ZAP/IFIT3/IFIT1-targeted Huh7.5 cells with the alphavirus Sindbis virus (SINV) or the unrelated rhabdovirus vesicular stomatitis virus (VSV) (Fig 4B). ZAP, IFIT3, and IFIT1 had modest but statistically significant contributions to the IFN-mediated suppression of both SINV and VSV. Together, our data suggest that IFN-mediated restriction of RNA viruses may be: 1.) completely independent of ZAP, IFIT3, and IFIT1, 2.) largely independent of ZAP, IFIT3, IFIT1, which play minor roles in viral restriction, 3.) largely dependent on ZAP, IFIT3, and IFIT1.

**Figure 4.**
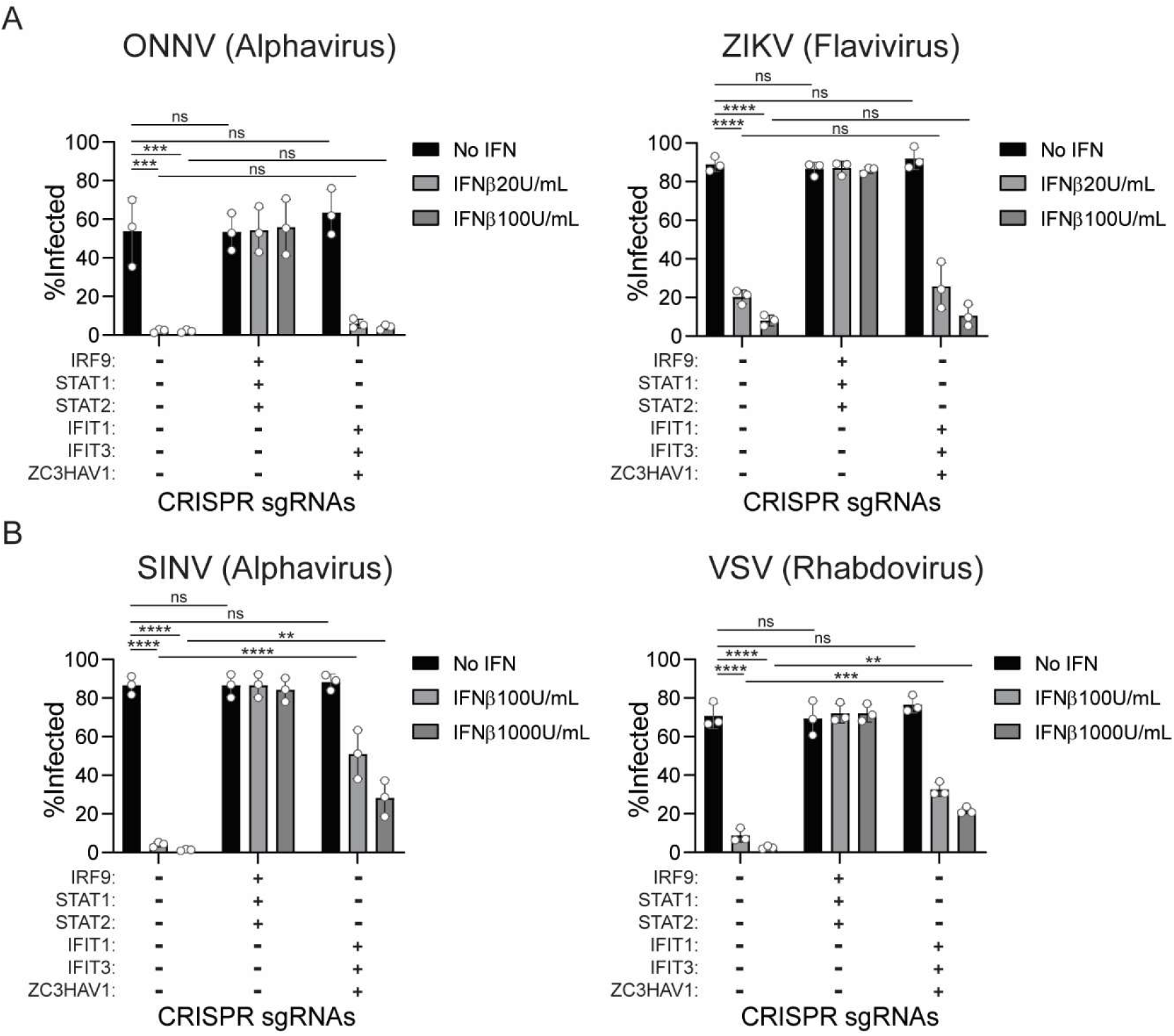
ZAP, IFIT3, and IFIT1 in combination exhibit differential antiviral activity towards diverse RNA viruses. A. U-2 OS control, IRF9/STAT1/STAT2-targeted, and ZAP/IFIT3/IFIT1-targeted cells were infected with GFP-expressing ONNV virus (left) or ZIKV (right) after IFNβ pretreatment. n=3, One-way AVONA with Dunnett’s test. B. Huh7.5 control, IRF9/STAT1/STAT2-targeted, and ZAP/IFIT3/IFIT1-targeted cells were infected with GFP-expressing Sindbis virus (AR86 strain, left) or vesicular stomatitis virus (right) after IFNβ pretreatment. The percentage of infected (GFP+) cells was quantified by flow cytometry. n=3, One-way ANOVA with Dunnett’s test.

The screens, validation, and triple CRISPR targeting experiments were evaluated by a single infection cycle of a VEEV with a virally encoded GFP reporter. We next sought to determine the role of ZAP, IFIT3, and IFIT1 in suppressing the production and spread of a non-reporter VEEV. To accomplish this, we infected control, ZAP/IFIT3/IFIT1-targeted cells or IRF9/STAT1/STAT2-targeted cells with non-reporter VEEV (TC-83 vaccine strain) after IFN pretreatment. We then collected supernatants containing VEEV particles after 12h (~3 replication cycles) and quantified viral titers in the supernatant by plaque assay (Fig 5A). In control cells expressing non-targeting sgRNAs, IFN suppressed VEEV production more than 3500-fold, and virus production was nearly completely restored in IRF9/STAT1/STAT2-targeted cells. In ZAP/IFIT3/IFIT1-targeted cells, IFN suppressed VEEV production by only 43-fold (compared to 3500-fold in non-targeting cells), indicating that the ZAP/IFIT3/IFIT1 combination comprises a major contribution to the IFN-induced suppression of VEEV production. Further demonstrating this, we observed an 86-fold increase in viral titers in IFN-treated ZAP/IFIT3/IFIT1-targeted cells relative to IFN-treated non-targeting cells. However, we still observed some IFN-mediated restriction after CRISPR targeting of ZC3HAV1, IFIT3, and IFIT1 in the context of viral production. This may be due to incomplete silencing of ZC3HAV1, IFIT3, and IFIT1 (Fig 5A, Western Blot and Fig S2A). Alternatively, this could be due to one or more antiviral effectors acting during assembly or egress, stages of the viral replication cycle that would not have played a role in the single-cycle VEEV-GFP infection assays. It is also possible that other IFN-induced genes that were not identified by our screens may play a role in restricting VEEV infection. Accordingly, we sought to determine if some of the previously described anti-alphavirus effectors contribute to this residual IFN-mediated restriction of VEEV (Jones *et al*, 2013; Li *et al*, 2013; Mahauad-Fernandez *et al*, 2014; Ooi *et al*, 2015; Poddar *et al*, 2016; Teng *et al*, 2012; Wan *et al*, 2019; Weiss *et al*, 2018; Weston *et al*, 2016; Zhang *et al*., 2007). We used multiplexed CRISPR-Cas9 to engineer U-2 OS cells that have up to 4 antiviral effectors simultaneously targeted. The ZAP/IFIT3/IFIT1-targeted cells were transduced with lentiCRISPv2 particles additionally targeting either *BST2* (which encodes Tetherin), *RSAD2* (which encodes Viperin), *IFITM3*, or *ISG20* (hereafter referred to as “quadruple-targeted” cells). After IFN pretreatment, we infected these quadruple-targeted cells with non-reporter VEEV and collected supernatants at 12h to quantify viral production. All of the quadruple-targeted cells produced similar levels of VEEV compared to the ZAP/IFIT3/IFIT1-targeted cells (Fig 5B, Fig S2B). This suggests that the residual IFN-mediated restriction in ZAP/IFIT3/IFIT1-targeted cells is not due to the previously described anti-alphavirus ISGs tetherin, viperin, IFITM3, or ISG20. The data further underscore the critical and dominant roles of ZAP, IFIT3, and IFIT1 in the IFN-mediated restriction of VEEV.

**Figure 5.**
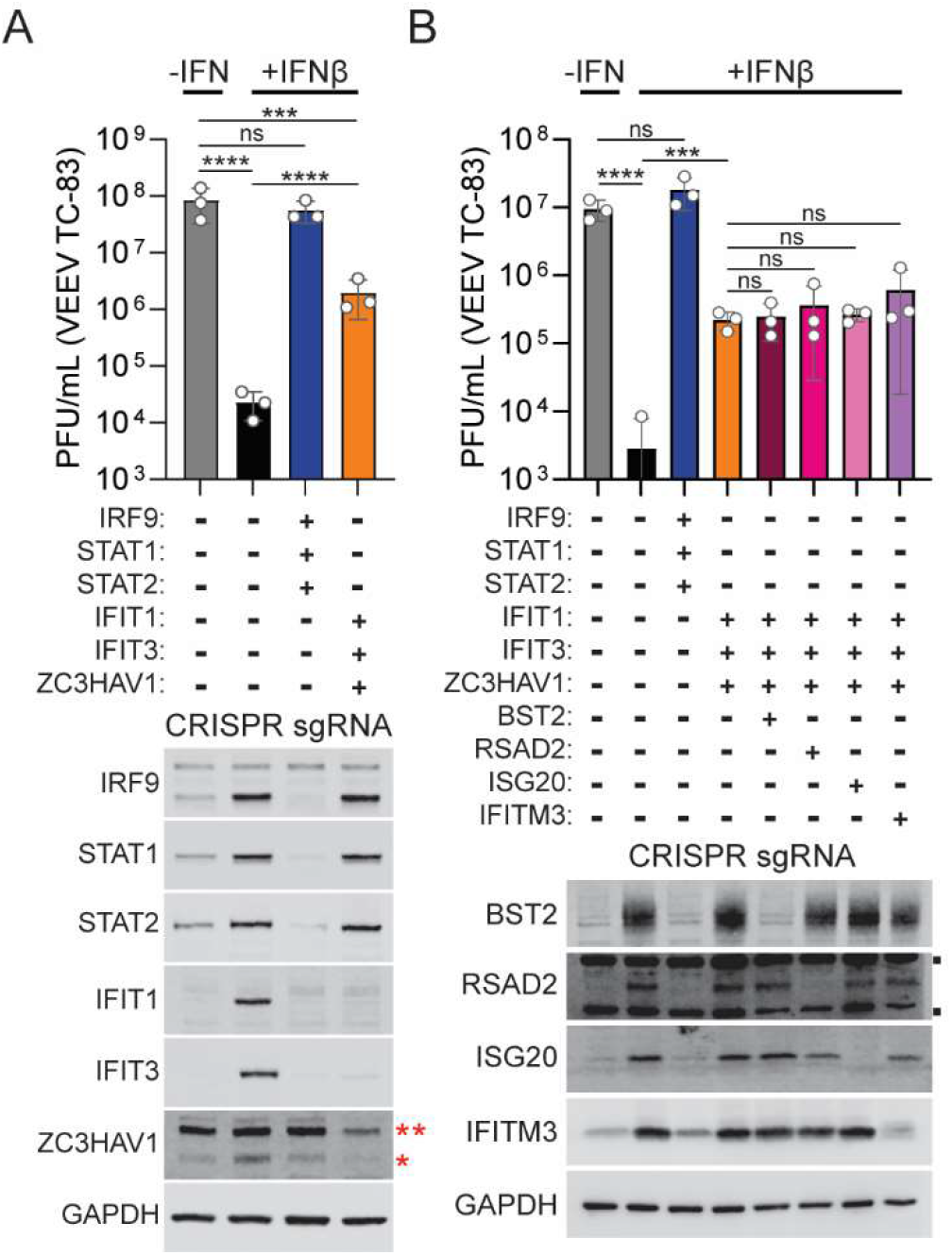
ZAP, IFIT3, and IFIT1, but not BST2, RSAD2, ISG20, or IFITM3 are required for the IFN-mediated suppression of VEEV production. A. (Top) Quantification of VEEV production (12 hrs post-infection) by plaque assay of control (far left: without IFN-pretreatment), IRF9/STAT1/STAT2-targeted, or ZAP/IFIT3/IFIT1-targeted cells after IFNβ pretreatment (IFNβ 20U/mL for 24 hrs). n=3, One-way ANOVA with Dunnett’s test on log-transformed data. (Bottom) Western blot shows IFN-induction and CRISPR targeting efficiency towards IRF9, STAT1, STAT2, ZAP, IFIT3, and IFIT1. Red asterisks denote long (**) and short (*) isoform of ZAP. B. (Top) Quantification of VEEV production (12 h post-infection) by plaque assay in control (far left: without IFN-pretreatment), IRF9/STAT1/STAT2-targeted, ZAP/IFIT3/IFIT1-targeted, or ZAP/IFIT3/IFIT1-targeted cells that were subsequently CRISPR targeted with a guide targeting BST2, RSAD2, ISG20, or IFITM3 after IFNβ pretreatment (IFNβ 20U/mL for 24 hrs). n=3, One-way ANOVA with Dunnett’s test on log transformed data. (Bottom) Western blot shows IFN-induction and CRISPR targeting efficiency towards BST2, RSAD2, ISG20, and IFITM3. Black squares denote non-specific bands.

Using CRISPR-Cas9 knockout screens followed by combinatorial ISG silencing, our study sheds light on the decades-old question of “how many ISGs are required for IFN-mediated protection of a particular virus?” (Staeheli *et al*., 1986). The data support a model in which the IFN-mediated suppression of VEEV is dominated by just three ISGs out of more than 600 IFN-inducible mRNAs (0.48% of the IFN-induced transcripts). Other ‘minor’ ISGs likely comprise the residual antiviral effect not accounted for by these three effectors. Notably, related alphaviruses like ONNV may to be sensitive to a distinct collection of ISGs, as ONNV was still inhibited by IFN in ZAP/IFIT3/IFIT1-targeted cells. We anticipate that ONNV may also be targeted by a limited number of ISGs, and that this hypothesis may generally extend to other diverse viruses. For example, in a previous CRISPR/ISG screen with the flavivirus, yellow fever virus, we discovered that the vast majority of IFN-mediated effects were surprisingly dominated by just one ISG, IFI6 (Richardson *et al*., 2018). In other CRISPR/ISG screens, unexpectedly small numbers of ISGs were also found to control HIV-1 and SARS-CoV-2 (Mac Kain *et al*., 2022; OhAinle *et al*., 2018). However, none of the CRISPR-based studies prior to the work presented here included combinatorial gene silencing to examine how many of the identified ISGs are needed for the IFN response. Nonetheless, from those studies and our current data, a pattern is emerging – that the IFN response to a particular virus may be governed by far fewer ISGs than previously anticipated. We speculate that certain ISGs will dominate an antiviral IFN response to a given virus, while others play more minor roles. This concept may be particularly important in studying polymorphisms of ISGs, as mutations in certain dominant antiviral effectors may be a determinant of severe viral disease. In addition, it may be important to prioritize mechanistic studies on ISGs that appear to exhibit more dominant roles in restricting viral infection rather than those that play minor roles. How this model might translate to other vertebrate species is not yet clear. It is possible that the set of human ISGs inhibiting a virus like VEEV may be partially overlapping, yet not completely identical to the set of ISGs inhibiting the same virus in a different animal. This is because of evolutionary principles governing the IFN response (McDougal *et al*, 2022). Certain ISGs like IFIT1, MX1, and TRIM5 are rapidly evolving and exhibit species-specific antiviral properties (Daugherty *et al*, 2016; Mitchell *et al*, 2012; Stremlau *et al*, 2004). Depending on the virus, the limited set of ISGs targeting that virus may differ from host to host, and dominant ISGs might represent important barriers to viral tropism. Additional studies, specifically using combinatorial ISG silencing, will be important for establishing whether the “limited set” model presented here for VEEV will translate to a broader paradigm that applies to other viruses and other host species. In addition, our study and others used type I IFNs for screening and subsequent follow-up experiments (IFNα and β) (Mac Kain *et al*., 2022; OhAinle *et al*., 2018; Richardson *et al*., 2018). Therefore, additional studies may be needed to determine if this “limited set” model also applies to the antiviral properties of type II and type III IFNs, or if the dominant effectors inhibiting a virus might differ depending on the IFN used.

## Materials and Methods

### Cells

U-2 OS, Huh7.5, and 293T cells and all derivatives were maintained in “complete” DMEM (Gibco) supplemented with 10% FBS (Gibco) and 1× non-essential amino acids (NEAA; Gibco). All derived stable cells expressing selectable markers were grown in complete DMEM containing puromycin (Gibco; U-2 OS: 0.5μg/mL, Huh7.5: 4μg/mL), blasticidin (Gibco; U-2 OS: 10μg/mL, Huh7.5: 15μg/mL), G418/Geneticin (Gibco; U-2 OS: 1 mg/mL, Huh7.5: 1mg/mL), and/or hygromycin (Invitrogen; U-2 OS: 250μg/mL, Huh7.5: N/A). BHK-21J cells were propagated in “complete” MEM (Gibco) supplemented with 10% FBS (Gibco) and 1x NEAA (Gibco). All cells were incubated at 37°C in 5% CO_2_.

### Viruses and viral infections

Venezuelan equine encephalitis virus (TC-83 strain) infectious clone was obtained from I.Frolov. A non-reporter VEEV infectious clone was subsequently generated using Gibson cloning to remove the GFP reporter. VEEV and VEEV-GFP (Petrakova *et al*, 2005), Sindbis virus (SINV-GFP, clone S300 from M. Heise) (Simmons *et al*, 2010), O’nyong ‘nyong virus (ONNV-GFP) (from S. Higgs) (Brault *et al*, 2004), and Zika virus (MR766-Venus, from M. Evans) (Schwarz *et al*, 2016) were all generated as previously described. Vesicular stomatitis virus (VSV-GFP) (from J. Rose) was propagated by passaging in BHK-21J cells and storing clarified supernatants as virus stocks at −80°C. All viral infections were performed in DMEM supplemented with 1% FBS and 1x non-essential amino acids at a total volume of 0.2mL in 24-well tissue culture plates. After a 1 h incubation period, 0.3mL of complete DMEM was added to all wells until harvest. For analysis of viral infection by flow cytometry, cells were dissociated from the plate with 150μL Accumax (Innovative Cell Technologies) and then fixed in paraformaldehyde at a final concentration of 1%. Cells were allowed to fix for at least 30 min at 4°C and subsequently centrifuged at 1500 x *g* for 5 min. Supernatant was removed from cell pellets and fixed cells were then resuspended in FACS buffer (PBS [Gibco] supplemented with 3% FBS). Flow cytometry was performed on the Stratedigm S1000EON benchtop flow cytometer, with analysis in FlowJo software.

IFN treatments were performed using human IFN Beta 1a (PBL Assay Science, Stock concentration of 6×10^5^ U/mL). Briefly, IFNβ was diluted to the indicated concentration in complete DMEM to a total volume of 0.5mL and incubated with cells for 24 h prior to infection.

### Genome-wide and ISG-targeted CRISPR screens

The human “Brunello” CRISPR knockout pooled library (#73179, Addgene) (Doench *et al*., 2016) and Human Interferon-Stimulated Gene CRISPR Knockout pooled Library (#125753, Addgene) (OhAinle *et al*., 2018; Roesch *et al*., 2018) were obtained from Addgene and amplified according to provided instructions. The genome-wide CRISPR screen was performed as 2 independent biological replicates at approximately 417x and 460x library coverage. The ISG-targeted CRISPR screen was performed as 3 independent biological replicates at approximately 1270x, 430x, and 1160x library coverage. The screening protocol was modified for U-2 OS cells and Venezuelan equine encephalitis virus infection from methods previously described (Richardson *et al*., 2018). Briefly, U-2 OS cells were transduced with a pooled CRISPR knockout library of lentivirus at an MOI between 0.5 and 1 for 48 hours. All cells were then selected for successful transduction by splitting into complete DMEM containing 1μg/mL of puromycin for 3 days and subsequently maintained in complete DMEM containing 0.5μg/mL of puromycin until plating. To perform the screen, transduced U-2 OS cells were plated and treated with IFNβ 20U/mL for 24 h prior to infection. Cells were then challenged with VEEV-GFP (~MOI 10) for 5 h, harvested, pooled, centrifuged, and resuspended in PBS containing 2% FBS and 0.5 mM EDTA for FACS. FACS was performed using a FACSAria II (Becton Dickenson) flow cytometer kept at 4°C to collect all GFP-positive cells (in PBS, 50% FBS, 50 mM HEPES). Sorted cells were then lysed and genomic DNA was extracted. PCR was then performed to amplify and barcode sgRNA sequences for next generation sequencing (NGS). 100μl PCR reactions (containing 6μg [Genome-wide screen] or 3μg of gDNA [ISG-targeted screen], Ex Taq polymerase, pooled P5 and barcoded P7 primers) were performed for each condition, and pooled. PCR product was purified for NGS using AMPure XP beads (Agencourt). Samples were then sequenced using Illumina NextSeq 500 and raw sequencing results were processed and mapped to the Brunello or Interferon-Stimulated Gene CRISPR Knockout Library. MAGeCK software was used for determining enriched sgRNAs in screening samples.

### Plaque Assays

For assessing viral production by plaque assay, after a 1h incubation of inoculum, cells were washed four times with 1mL of 10% DMEM. Subsequently, 0.5mL of complete DMEM was added to cells and collected 12 h post-infection and stored at −80°C. Plaque assays were then performed on BHK-21J cells. Briefly, BHK-21J cells were plated at 450,000 cells per well in 6-well plates. Thawed supernatants were then diluted in 1% MEM, and 200μL of diluted supernatants were applied to cells. Plates were incubated for 1h at 37°C with intermittent rocking every 10-15min, after which 2mL of semi-solid overlay (1x DMEM, 1X Pen/Strep, 4% FBS, 10mM HEPES, 1.2% Avicel, 0.1125% NaHCO_3_) was added to all wells. After 48h, cells were fixed with formaldehyde for at least 30 min and stained with crystal violet staining solution (1.28% crystal violet powder dissolved in 40% ethanol). Plaques were counted on relevant dilutions to calculate plaque forming units per mL (PFU/mL) of virus in supernatant.

### Cloning and plasmids

CRISPR sgRNA guides were cloned in the LentiCRISPRv2 plasmid backbone as previously described (Sanjana *et al*, 2014; Shalem *et al*, 2014). All oligo sequences used for generating CRISPR sgRNAs were obtained from the Brunello library. All LentiCRISPRv2 plasmids (expressing a selectable marker of resistance to puromycin, blasticidin, neomycin, or hygromycin) were purchased from Addgene and amplified according to instructions. LentiCRISPRv2 (puro) was a gift from Feng Zhang (Addgene plasmid # 52961). LentiCRISPRv2 hygro, LentiCRISPRv2 neo, and LentiCRISPRv2 blast were gifts from Brett Stringer (Addgene plasmids # 98291, 98292, 98293).

### Western Blotting

Cells were lysed with M-PER (Mammalian Protein Extraction Reagent, Thermo Scientific) containing 1x complete protease inhibitor cocktail (Roche) at 4°C for 5 min. Lysate was then stored at −80°C. For running SDS-PAGE, lysate was thawed on ice, and mixed to a final concentration of 1x SDS Loading Buffer (0.2 M Tris-Cl pH 6.8, 5% SDS, 25% Glycerol, 0.025% Bromophenol Blue, and 6.25% beta-mercaptoethanol). Samples were run on 12% TGX FastCast acrylamide gels (Bio-Rad) and transferred to nitrocellulose membranes using a TransBlot Turbo system (Bio-Rad). Membranes were blocked with 5% milk in 1x Tris Buffered Saline, with Tween-20 (TBS-T [20mM Tris, 150mM NaCl, 0.1% Tween-20]) for at least 45min. Subsequent incubation with primary antibody (at dilutions between 1:500 to 1:5000) diluted in 1% milk in TBS-T was performed overnight, rocking at 4°C. After incubation with primary antibodies, membranes were washed 3 times with TBS-T by rocking for 5 min. Secondary antibodies (Goat anti-Rabbit 800CW and Goat anti-Mouse 680RD [LI-COR] at 1:5000 dilutions) diluted in 1x TBS-T were incubated with membranes for at least 60 min. Membranes were then washed 3 times with TBS-T by rocking for 5 min. Proteins were visualized using the LI-COR Odyssey FC imaging system.

### Lentivirus production and transduction

To generate lentivirus pseudoparticles, 350,000 293Ts were plated onto poly-lysine coated 6-well plates. Lentiviral packaging plasmids expressing VSV-g and Gag-pol were then mixed with LentiCRISPRv2 plasmids at a ratio of 0.2μg:0.8 μg:1 μg, respectively. Plasmids were incubated for 20 min with a mix of XtremeGENE9 transfection reagent (Roche) and OptiMEM (Gibco). Transfection complexes were then added to 293T cells in 1mL of DMEM supplemented with 3% FBS dropwise. After 6 h of incubation, media was changed to 1.5 mL of DMEM supplemented with 3% FBS. Supernatants containing lentiviral pseudoparticles were harvested at 48 and 72 h post-transfection. Lentiviral pseudoparticles were then processed by adding 4μg/mL Polybrene (Sigma), 20mM HEPES, centrifuging at 1500rpm for 5min to remove cell debris, and storing at - 80°C until transductions.

Lentivirus transductions were performed in 6-well plates after plating 200,000 cells (U-2 OS) or 350,000 cells (Huh7.5). Final volume of transductions was either 1mL or 1.5mL of lentivirus mixed with “pseudoparticle” DMEM (DMEM supplemented with 3% FBS, 4μg/mL Polybrene, and 20mM HEPES) After a 6 h incubation at 37°C, 2mL of complete DMEM was added to all transductions. Transduced cells were split into selection media 48h post-transduction and passaged for a minimum of three passages in selection media.

### RNA-Sequencing to determine IFN-induced transcripts

To evaluate for transcripts upregulated by IFN, 300,000 U-2 OS cells were plated in 6-well plates and treated the following day with complete DMEM containing 20U/mL of IFNβ (PBL Assay Science). RNA was harvested at 6 and 24 h post IFN-treatment (three independent biological triplicates) and isolated using the RNeasy Mini Kit (Qiagen). RNA-Seq was performed by Novogene using a PE150 sequencing strategy on the Illumina NovaSeq 6000 platform. Briefly, 1 μg RNA per sample was used as input and mRNA was enriched using poly-T oligo-attached magnetic beads. Libraries were generated using NEBNext UltraTM RNA Library Prep Kit for Illumina (NEB) following manufacturer’s recommendations. Sequencing reads were then mapped to the human genome using the Spliced Transcripts Alignment to a Reference (STAR) software. DESeq2 (Love *et al*, 2014) was then used to determine genes that were differentially expressed after IFN treatment. P-values were adjusted using Benjamini-Hochberg method for controlling the False Discovery Rate (FDR). Level of gene expression was determined by using HTSeq v0.6.1 software to count the reads mapped to each gene. Fragments Per Kilobase of transcript per Million mapped reads (FPKM) of each gene was calculated based on gene length and read counts mapped to the corresponding gene (Mortazavi *et al*, 2008).

### Statistical analysis

All statistical analyses were performed in GraphPad Prism 9. All graphs represent the mean of n=3 independent biological replicates with error bars representing standard deviation. Statistical analysis for CRISPR screening was done with MAGeCK analysis. Statistical analysis for RNA-Seq was performed with DESeq2. ns, p > 0.05, *, p < 0.05, **, p < 0.01, ***, p < 0.0001, and **** p < 0.0001.

## Acknowledgements

We would like to thank Julie Pfeiffer for critical reading of the manuscript. This work was funded in part by the NIH Grant AI158124 (J.W.S), the Clayton Foundation for Research (J.W.S.), the Welch Foundation (20190330 to J.W.S), and the National Science Foundation Graduate Research Fellowship (2019274212 to M.B.M.). John W. Schoggins holds an Investigators in the Pathogenesis of Infectious Disease Award from the Burroughs Wellcome Fund.

## Supplementary Figures

**Figure S1.**
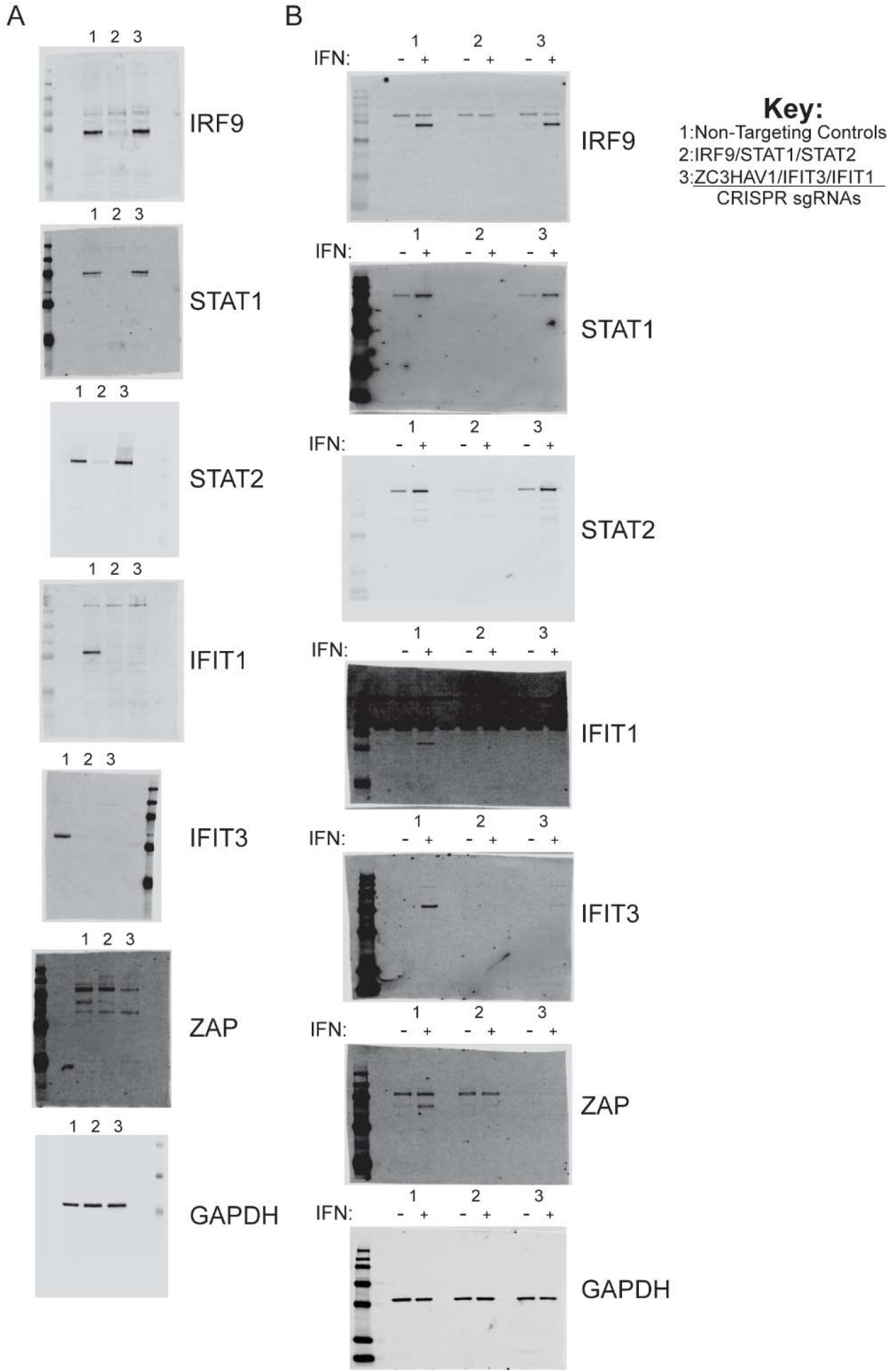
Full membranes from western blots demonstrating CRISPR targeting efficiency in U-2 OS and Huh7.5 cells. A. Full membranes from Western blot showing CRISPR targeting efficiency at the protein-level in control, IRF9/STAT1/STAT2-targeted, or ZAP/IFIT3/IFIT1-targeted U-2 OS cells after IFNβ pretreatment (IFNβ 20U/ml for 24 hrs) (See Fig 3A for cropped membranes). Blots represent one of three independent biological replicates showing similar results. B. Full membranes from Western blot showing CRISPR targeting efficiency at the protein-level in control, IRF9/STAT1/STAT2-targeted, or ZAP/IFIT3/IFIT1-targeted Huh7.5 cells with and without IFNβ pretreatment (IFNβ 100U/ml for 24 hrs) (See Fig 3B for cropped membranes). Blots represent one of three independent biological replicates showing similar results.

**Figure S2.**
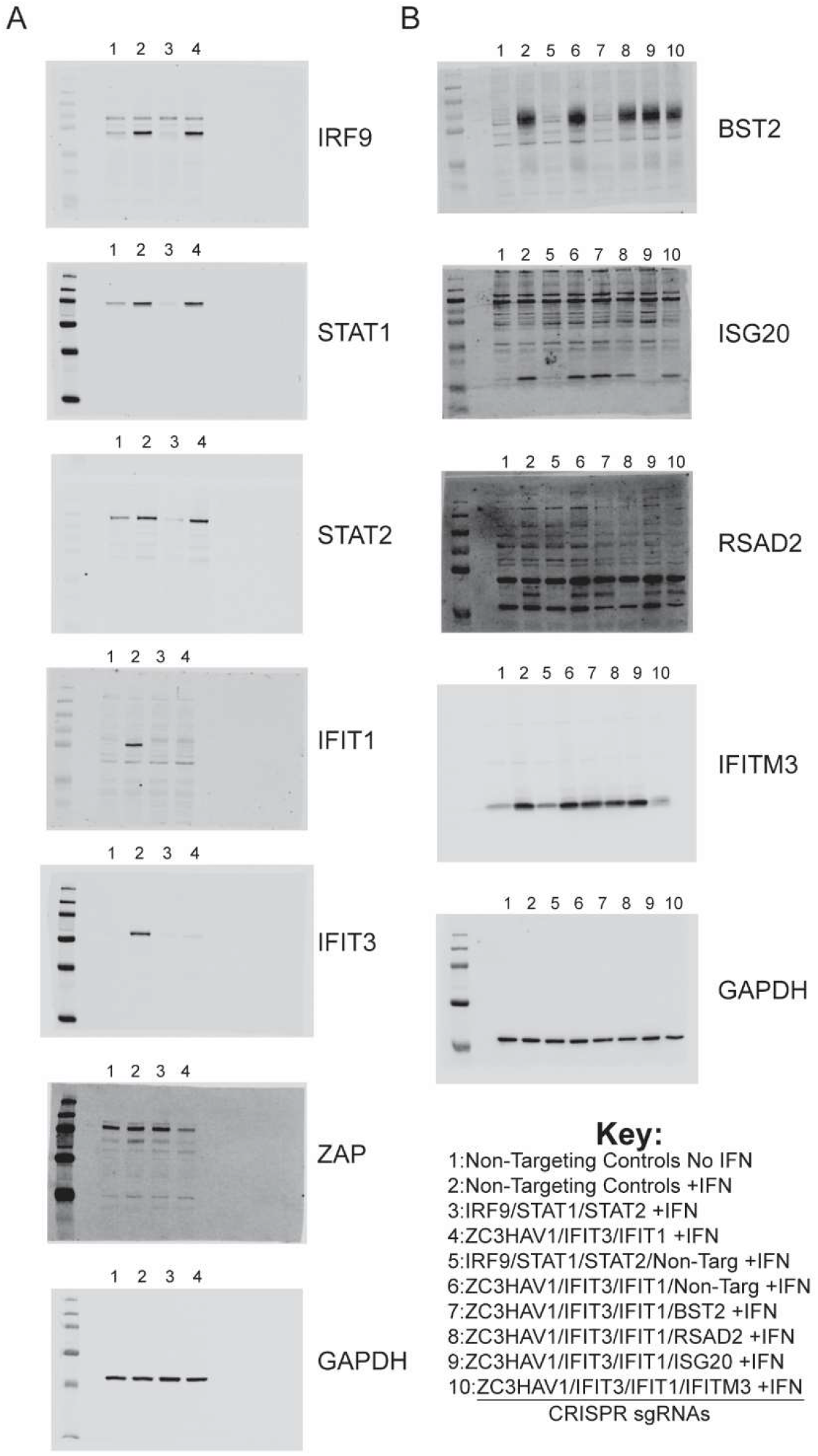
Full membranes from western blots demonstrating CRISPR targeting efficiency in U-2 OS cells used for viral production assays. A. Full membranes from Western blot showing CRISPR targeting efficiency at the protein-level in control (far left: without IFN-pretreatment), IRF9/STAT1/STAT2-targeted, or ZAP/IFIT3/IFIT1-targeted U-2 OS cells after IFNβ pretreatment (IFNβ 20U/ml for 24 hrs) (See Fig 5A for cropped membranes). Blots represent one of three independent biological replicates showing similar results. B. Full membranes from Western blot showing IFN-induction and CRIS PR-targeting efficiency towards BST2, RSAD2, ISG20, and IFITM3 at the protein-level in "quadruple-targeted" U-2 OS cells with or without (far left) IFNβ pretreatment (IFNβ 20U/ml for 24 hrs) (See Fig 5B for cropped membranes). Blots represent one of three independent biological replicates showing similar results.

